# Alpha-1-antitrypsin (AAT) inhibits *Mycobacterium intracellulare* induction of monocyte colony stimulating factor: another host-defense function of AAT

**DOI:** 10.64898/2026.02.20.706890

**Authors:** Xiyuan Bai, Drew E. Narum, Carla V. Eyre, Junfeng Gao, Edward D. Chan

**Affiliations:** Department of Medicine, Rocky Mountain Regional Veterans Affairs Medical Center, Aurora, CO, USA; Department of Medicine, National Jewish Health, Denver, CO, USA; Department of Academic Affairs, National Jewish Health, Denver, CO, USA; Department of Immunology and Genomic Medicine, National Jewish Health, Denver, CO, USA; Department of Biostatistics and Informatics, University of Colorado School of Public Health Anschutz Medical Campus, Aurora, CO, USA; Division of Pulmonary Sciences and Critical Care Medicine, University of Colorado Anschutz Medical Campus, Aurora, CO, USA

**Keywords:** serine protease, gene regulation, AAT, glucocorticoid receptor, GM-CSF

## Abstract

**RATIONALE:** The host-protective role of alpha-1-antitrypsin (AAT) against mycobacteria may be partly attributed to its binding to the cytoplasmic glucocorticoid receptor (GR) that results in gene regulation in macrophages that favors killing of ingested mycobacteria. The AAT–GR complex was found to be significantly responsible for limiting *Mycobacterium avium* complex (*MAC*) burden in macrophages; this host-protective function of AAT–GR is due, in part, to induction of *COLONY STIMULATING FACTOR-2* (*CSF-2*) gene which encodes for granulocyte-monocyte colony stimulating factor (GM-CSF).

**METHODS:** To better understand the role of AAT–GR binding during mycobacterial infection, we performed bulk RNA sequencing (RNA-seq) on four different groups of cells: *(i)* control THP-1 cells (THP-1^control^); *(ii)* THP-1^control^ cells infected with *Mycobacterium intracellulare* (*MAC*); *(iii)* THP-1^control^ cells incubated with *MAC* + AAT; and *(iv)* THP-1 cells knocked down for GR (THP-1^GR-KD^) incubated with *MAC* + AAT.

**RESULTS:** Our analyses revealed that *MAC* infection significantly upregulated 1,977 genes and significantly downregulated 2,303 genes in THP-1^control^ cells. Additionally, AAT significantly upregulated 1,200 genes and downregulated 890 genes in *MAC*-infected THP-1^control^ cells. Furthermore, the regulation of 1,624 genes that are regulated by AAT+GR in THP-1^control^ cells was augmented in THP-1^GR-KD^ cells, indicating that the regulation of these genes by AAT+*MAC* is inhibited by GR. Conversely, the regulation of 1,683 genes by AAT+*MAC* in THP-1^control^ cells was attenuated in THP-1^GR-KD^ cells, indicating that the regulation of these genes by AAT+*MAC* is enhanced by GR. *MAC* also induced both *CSF2* (GM-CSF) and *CSF1* (encodes for monocyte colony stimulating factor, M-CSF) expression. Whereas AAT inhibited *MAC*-induced M-CSF mRNA was dependent on GR, this inhibition of M-CSF protein was not dependent on GR. In contrast, AAT did not inhibit *MAC*-induced *CSF2* (GM-CSF) expression. Since either *MAC* or AAT induced GM-CSF expression in macrophages, further investigation revealed that AAT-inhibition of cell-associated *MAC* burden was abrogated upon neutralization of endogenous GM-CSF.

**CONCLUSIONS:** The ability of AAT to induce GM-CSF and to inhibit *MAC*-induced M-CSF may skew macrophages to a phenotype that is better endowed to control mycobacterial infection.

## INTRODUCTION

Non-tuberculous mycobacteria (NTM) infections in humans manifest as chronic lung disease, lymph node infections, skin/soft tissue/osteoarticular infections, or disseminated disease [1, 2]. Since the prevalence and mortality of NTM lung disease have increased in the past two decades [3–8], due in large part to a high level of intrinsic drug resistance, it is imperative to develop more effective antimicrobials and host-directed treatments.

Alpha-1-antitrypsin (AAT) is the most prevalent serine protease inhibitor (serpin) in tissues and bloodstream, with normal circulating levels in humans of 100-200 mg/dL. The canonical function of AAT is the binding and inhibition of elastase and other serine proteases [9–12]. AAT also has anti-inflammatory and host-defense properties independent of its serpin function [13–17]. We demonstrated that AAT enhances both phagosome-lysosome fusion and autophagic flux – mechanisms used by macrophages to control infection with *Mycobacterium intracellulare*, a species within the *Mycobacterium avium* complex (*MAC*) [13]. While the exact mechanisms by which AAT affects these host-protective functions are varied and complex, evidence suggests involvement of gene regulation [13, 14, 18].

We reported that AAT binds to the glucocorticoid receptor (GR) to regulate gene expression [14, 18]. The GR normally resides in the cytoplasm in a sequestered inactive state. Upon binding to glucocorticoids, its canonical ligand, the glucocorticoid–GR complex translocates into the nucleus to induce or inhibit the expression of a vast array of genes [19–21]. Analogously, we found that the binding of AAT to GR in macrophages displayed anti-inflammatory and host-protective effects. This interaction results in inhibition of lipopolysaccharide-induced nuclear factor-kappa B (NFκB) activation and interleukin-8 (IL-8) production [14]. Additionally, the intracellular burden of *Mycobacterium tuberculosis* and *M. intracellulare* (which we will henceforth refer to as *MAC*) was reduced by the AAT–GR complex. Subsequently, we reported an expansive array of genes that are either inhibited or induced by AAT in a GR-dependent manner, many of which have been shown to predispose or protect against mycobacterial infections [18]. One gene that was shown to be strongly induced by AAT (20-fold) in a GR-dependent manner is *Colony Stimulating Factor-2* (*CSF2*), which encodes for granulocyte-monocyte colony stimulating factor (GM-CSF) [18].

To determine the role of the AAT–GR complex with mycobacterial infection, we performed bulk RNA sequencing (RNA-seq) on control THP-1 macrophages and THP-1 macrophages stably knocked down for GR under conditions of *MAC* infection and addition of exogenous AAT. Our analyses revealed that *MAC* infection of macrophages induced the expression of both *CSF2* and *CSF1* genes, encoding for GM-CSF and monocyte-colony stimulating factor (M-CSF), respectively. While AAT inhibited *MAC*-induced M-CSF mRNA expression in a GR-dependent fashion, AAT inhibition of *MAC*-induced M-CSF protein was observed to be independent of GR, suggesting that AAT may have GR-independent effects post-transcriptionally. Whereas GM-CSF differentiates macrophages to the M1 phenotype, characterized by a greater capacity to kill intracellular pathogens, M-CSF skews macrophages to the M2 phenotype. M2 macrophages have less capacity to kill intracellular pathogens but are nevertheless important in dampening any excessive and injurious immune activation during the resolution phase of an infection. These findings elucidate a previously unknown mechanism by which AAT plays a host-protective role against mycobacterial infections.

## MATERIAL and EXPERIMENTAL METHODS

### Materials

The human monocytic cell line (THP-1) was obtained from the American Type Culture Collection (Manassas, Virginia). Phorbol myristate acetate (PMA) was purchased from Sigma (St. Louis, MO). *Mycobacterium intracellulare* 9141 was obtained from the Clinical Mycobacterial Laboratory at National Jewish Health, Denver, Colorado. Fetal bovine serum (FBS) was purchased from Atlanta Biologicals (Lawrenceville, GA) and heat-inactivated at 56°C for 30 minutes. An aliquot of the glucocorticoid receptor (*NR3C1*) human shRNA (shRNA-GR) Lentiviral Particle was a kind gift from Miles Pufall, Ph.D. (University of Iowa Carver College of Medicine). RPMI-1640 and ELISA kit for human macrophage-colony stimulating factor (M-CSF) (#EHCSF1) were purchased from ThermoFisher Scientific (Carlsbad, CA). RNeasy Plus Mini kit for the purification of total RNA from cells was purchased from QIAGEN (Redwood City. CA). AAT (Glassia^®^) (NDC 0944-2884-01) was acquired from Kamada Ltd., Israel. The RT-qPCR primers were synthesized from Integrated DNA Technologies (Coralville, IA). The cDNA was synthesized using M-MLV Reverse Transcriptase kit from Promega (Fitchburg, WI). SYBR^TM^ Green PCR Master MIX were obtained from Applied Biosystems (ThermoFisher Scientific, #4309155).

### Differentiation, stable knockdown of the glucocorticoid receptor (GR) and *M. intracellulare* infection of THP-1 cells

Human THP-1 cells were cultured in RPMI-1640 medium containing 2 mM L-glutamine (Gibco; Grand Island, NY), 10% FBS, penicillin (100 U/mL), and streptomycin (100 µg/mL) at 37°C and 5% CO_2_. THP-1 cells were differentiated into macrophages following incubation with 15 ng/mL PMA for 24 hours. We employed shRNA-lentivirus technology to develop a pool of THP-1 cells stably knocked down (KD) for GR (THP-1^GR-KD^) and control THP-1 cells (THP-1^control^) using shRNA-GR-lentivirus and shRNA-scrambled-lentivirus, respectively, as previously described [14]. Following differentiation, the THP-1^GR-KD^ cells were confirmed to be depleted of GR mRNA and protein through RNAseq and immune-blotting, respectively [14]. Then the THP-1^control^ were left uninfected, infected with *MAC* at a multiplicity-of-infection (MOI) of 10 mycobacteria:1 macrophage, and infected with *MAC* plus incubated with AAT (3 mg/mL). In addition, THP-1^GR-KD^ were incubated with *MAC* (MOI 10) + AAT (3 mg/mL). After one hour of infection, the cells were washed to remove any uningested *M. intracellulare*, and incubated with replenished AAT (3 mg/mL) for an additional 48 hours. In essence, four experimental conditions comprise this study: *(i)* control THP-1 cells (THP-1 ^control^); (*ii*) THP-1^control^ cells infected with *MAC*; *(iii)* THP-1 ^control^ cells incubated with both *MAC* and AAT; and *(iv)* THP-1 cells knocked down for GR (THP-1 ^GR-KD^) incubated with both *MAC* and AAT.

### RNA isolation and RNA sequencing (RNA-seq)

For each of the four conditions, high-quality total RNA (260 nm/280 nm ∼ 2) was isolated from six million plated cells using RNeasy Plus kit per manufacturer’s instruction. The RNA-seq libraries prepared from independent experiments were prepared as described [22]. Total RNA from two independent experiments was sequenced using the Illumina NovaSeq 6000 from Genomics Shared Resource at the University of Colorado Cancer Center.

### RNA-seq data analysis

The RNA-seq data were analyzed as described previously [22]. Briefly, the raw reads (average 20 million paired end reads, two biological replicates for each treatment) were analyzed and quality checked by FastQC. The reads were aligned to the hg38 reference genome using the Spliced Transcripts Alignment to a Reference (STAR, version 2.4.0.1) software. Reads (FPKM) were assembled into reference transcripts and counted using Cufflinks (version 2.2.1). The average reads from two biological samples were calculated using Cuffmerge (version 1.0.0). The differential gene expression between the resting and stimulated samples was analyzed using Cuffdiff (version 2.2.1). Gene Ontology (GO) enrichment analysis was performed on AAT-regulated genes by using the Database for Annotation, Visualization and Integrated Discovery (DAVID, version 6.8) [23, 24]. The fold change of the genes was calculated using the following ratios: MAC infected / uninfected THP-1^control^ cells, AAT-stimulated/unstimulated MAC-infected THP-1^control^ cells, and MAC+AAT-stimulated THP-1^GR-KD^/MAC+AAT-stimulated THP-1^control^ cells. The genes with a fold change ≥2 were classified as “highly upregulated” while the genes with a fold change ≤0.5 were classified as “highly downregulated.” Thus, AAT-stimulated genes in which THP-1^GR-KD^ /THP-1^control^ ≤0.5 would be those that are dependent on GR. Several genes relevant in mycobacterial infection and overall host immunity were selected from each group for heatmap representations, which were generated using Expander software (version 7.2) [25].

### Reverse transcription and quantitative polymerase chain reaction (RT-qPCR)

Following incubation with medium alone, *MAC* infection, or both AAT + *MAC* of THP-1^control^ macrophages as well as incubation with AAT + *MAC* of THP-1^GR-KD^ macrophages for 48 hours, total RNA was isolated. The cDNA was then prepared and quantitative PCR was performed in a QuantStudio 7 Flex Real-Time PCR System using SYBR^TM^ Green qPCR Master MIX. Relative amount of mRNA = 2^[Ct(*Sample*)-Ct(*HPRT*)]^ where *HPRT* is the housekeeping gene that encodes for hypoxanthine phosphoribosyltransferase 1.

### ELISA

After being incubated with medium alone or with AAT (3 mg/mL) for 48 hours, the supernatants of differentiated THP-1^control^ or THP-1^GR-KD^ macrophages with or without *MAC* infection were quantified via ELISA for M-CSF following manufacturer’s instruction.

### Quantitation of cell-associated *MAC* infection

THP-1 cells were cultured in RPMI-1640 medium containing 10% FBS and 100 U/mL penicillin, 100 µg/mL streptomycin at 37°C and 5% CO_2_. The cells were differentiated into macrophages following incubation with 15 ng/mL of PMA for 24 hours. After washing with wash buffer (1:1 of RPMI-1640 medium:PBS), the differentiated macrophages were infected with *MAC* at an MOI of 10 mycobacteria:1 macrophage. After one hour, two and four days of incubation, the cells were washed, lysed, the lysates serially diluted, plated on 7H10 solid medium, and cell-associated *MAC* quantified as previously described [14].

### Statistical Analysis

Replicate RNA-seq experiments are independent and presented as mean ± SD. The ELISA data are reported as mean ± SEM of duplicate samples of three independent experiments. Group means were compared by repeated-measures ANOVA using Fisher’s least significant test or by two-way ANOVA with Bonferroni’s post-hoc test. Data were graphed in Prism 9^®^ and comparisons were considered significant when p<0.05.

## RESULTS

### Genes regulated by *MAC* infection of THP-1^control^ macrophages

To determine the effect of *MAC* infection on gene expression of THP-1^control^ cells, unstimulated THP-1^control^ cells were left unstimulated or infected with *MAC*. After 48 hours, total RNA was isolated and sequenced. Gene ontology (GO) enrichment analysis was used to first categorize genes upregulated (**Supplemental Figure 1A**) or downregulated (**Supplemental Figure 1B**) by *MAC* infection of THP-1^control^ cells into various functional categories/pathways. Comparison of THP-1^control^ and THP-1^control^ + *MAC* revealed that *MAC* infection significantly upregulated 1,977 genes and downregulated 2,303 genes. The heat maps of specific genes upregulated or downregulated by *MAC* infections of THP-1^control^ cells were generated (**Figures 1 and 2**). Several genes relevant for mycobacterial infections that were upregulated by *MAC* include *IL23A, IL1B*, *CSF2*, *CCL3*, *TNF*, and *CSF1* (**Figure 1**). Relevant gene downregulated by *MAC* include *CX3CR1* and matrix metalloproteinases (*MMP11*, *MMP15*, *MMP16*, and *MMP17*) (**Figure 2**). We highlight some of the *MAC*-regulated genes with functional relevance for mycobacterial infections (**Table 1**).

**Figure 1.**
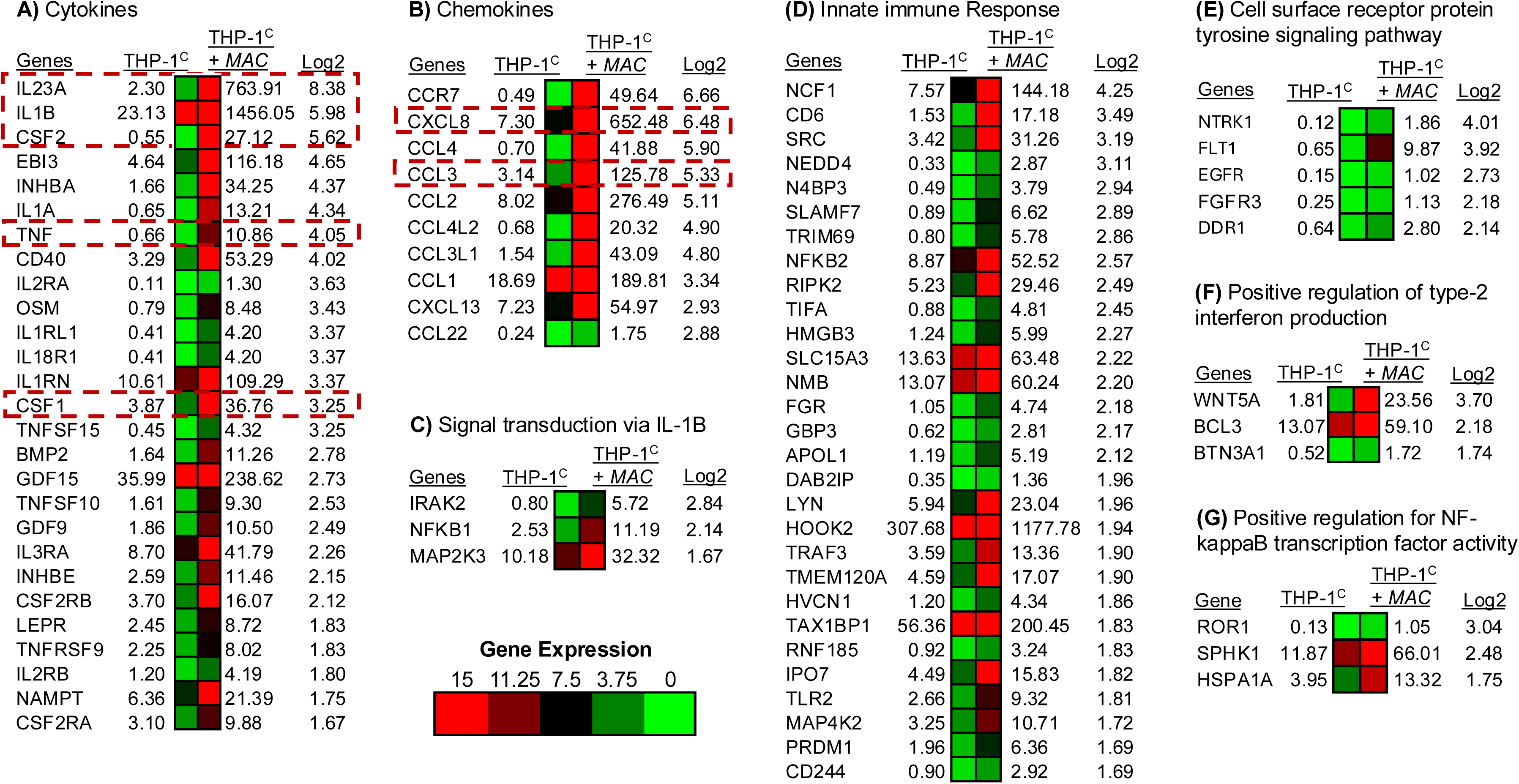
Heatmap of selected genes upregulated by *MAC* in THP-1^C^ macrophages. Comparison of gene expression in unstimulated vs. *MAC*-infected THP-1^C^ cells to determine what significant genes are upregulated during *MAC* infection. Significantly upregulated genes are defined by a change of ≥ two-fold (log_2_≥1) of the Reads per Kilobase Million (RPKM) of [THP-1^C^+*MAC*] divided by that of THP-1^C^ = numbers listed below log_2_. The heatmap is a visual representation of gene expression for THP-1^C^ and THP-1^C^+*MAC*. Some of the relevant genes in host defense and mycobacterial immunity were selected and sorted into the following functional categories: **(A)** Cytokines; **(B)** Chemokines; **(C)** Signal transduction through IL-1B; **(D)** Innate immune response; **(E)** Cell surface receptor protein tyrosine signaling pathway; **(F)** Positive regulation of type-2 interferon production; and **(G)** Positive regulation for NF-kappa B transcription factor activity. **IL-1B**=Interleukin-1 beta; ***MAC***=*Mycobacterium avium* complex; **THP-1^C^**=control THP-1 cells.

**Figure 2.**
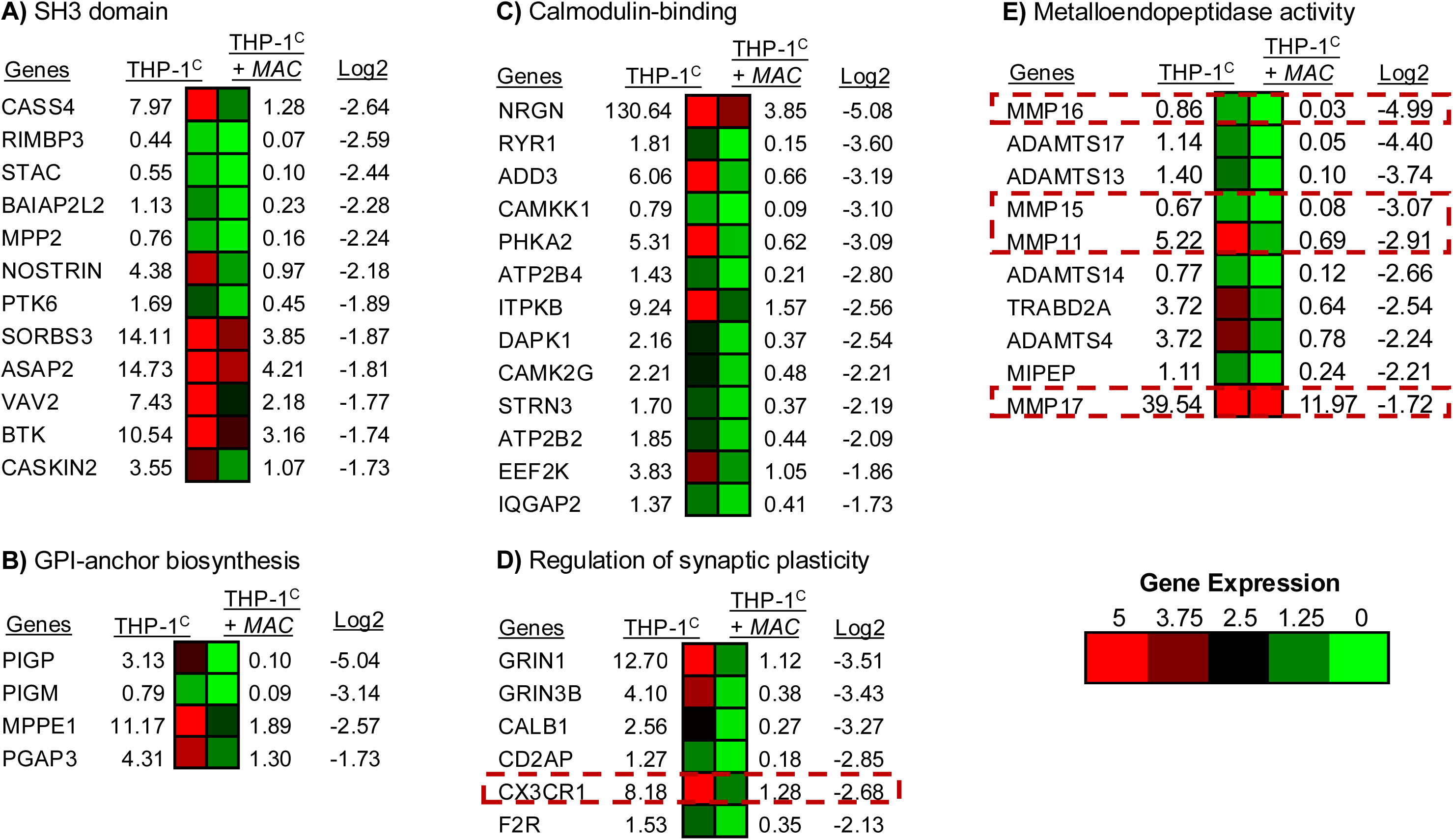
Heatmap of selected genes downregulated by *MAC* in THP-1^C^ macrophages. Comparison of gene expression in unstimulated vs. *MAC*-infected THP-1^C^ cells to determine what significant genes are downregulated during *MAC* infection. Significantly downregulated genes are signified by a change of ≤ 0.5-fold (log_2_≤–1) of the Reads per Kilobase Million (RPKM) of [THP-1^C^+*MAC*] divided by THP-1^C^ = numbers listed below log_2_. Some of the relevant genes in host defense and mycobacterial immunity were selected and sorted into the following functional categories: **(A)** Src homology 3 domain; **(B)** GPI-anchor biosynthesis; **(C)** Calmodulin-binding; **(D)** Regulation of synaptic plasticity; and **(E)** Metalloendopeptidase activity. ***MAC***=*Mycobacterium avium* complex; **THP-1^C^**=control THP-1 cells.

**Table 1.** *MAC*-regulated genes relevant for mycobacterial infections.

| Gene | Protein | Brief synopsis of function |
| --- | --- | --- |
| **Selected *MAC*-upregulated genes** |  |  |
| ***IL23A*** | Interleukin 23 alpha subunit (p19) | IL-23 is essential to host immunity against mycobacteria, as it is important for differentiation of T cells to the T<sub>H</sub>1 and T<sub>H</sub>17 phenotypes [26] as well as to IFNγ-dependent in Vδ2<sup>+</sup> $\gamma\delta$ T cells and MAIT cells [27, 28]. |
| ***CXCL8*** | CXCL8 (IL-8) | CXCL8 (interleukin-8) is a pro-inflammatory chemokine produced in response to mycobacterial infection. Mycobacterial components stimulate CXCL8 production through Toll-like receptor signaling [29, 30].<br>IL-8 is a chemokine necessary for recruitment and activation of neutrophils at the site of infection in the early innate immune response. Neutrophils contribute to initial containment of *M. tuberculosis*, but neutrophilia is also associated with increased mortality from TB [31, 32].<br>CXCL8-mediated recruitment is associated with lung inflammation and tissue damage [33]. CXCL8 is expressed within tuberculous granulomas and elevated levels correlate with severity of active disease, making it a potential biomarker of inflammation in tuberculosis [33].<br>IL-8 is secreted from neutrophils at a significantly higher rate by *MAC*-treated neutrophils than *M. gordonae* and *M. smegmatis* [34].<br>IL-8 mRNA is expressed more in THP-1 macrophages infected with *M. avium* than *M. abscessus* complex, *M. smegmatis*, and *M. tuberculosis* [35]. |
| ***IL1B*** | Interleukin-1β | Host-protective qualities of IL-1β in *M. tuberculosis* infection is controversial, as there is evidence for and against this [36].<br>IL-1β aids in host defense against *M. tuberculosis* by enhancing macrophages antimicrobial activity through upregulation of TNF and TNF receptor expression, promoting caspase-dependent killing of the mycobacteria [37]. |
|  |  | • Mutations in the *IL1B* gene in humans that result in increased IL-1β expression are associated with increased tuberculosis severity, potentially due to increased neutrophil infiltration [36, 38].<br>• In *M. kansasii* infection of THP-1 macrophages, blocking IL-1β with neutralizing antibody led to higher burden of intracellular bacteria [39].<br>• In mice infected with *M. abscessus*, IL-1β significantly contributed to lung inflammation and inflammatory IL-17 production [40]. It did not impact intracellular growth in macrophages and bacterial clearance in the lungs [40]. |
| **CSF2** | Granulocyte-monocyte colony stimulating factor (GM-CSF) | • Host-protective against *M. tuberculosis* and NTM [41-44], protects mice against *M. abscessus* infection [45].<br>• Induces macrophages towards "M1" phenotype which has greater capacity to dispose of intracellular pathogens, including MAC [46, 47].<br>• Augments control of *M. tuberculosis* in human macrophages, co-culture of invariant natural killer T (iNKT) cells and macrophages, and mice. GM-CSF works as a mediator in the CD1d signaling iNKT cells require [48].<br>• Necessary for surfactant recycling as lack of GM-CSF leads to surfactant accumulation in alveolar macrophages, impairing their function [49].<br>• GM-CSF modulates the inflammatory response through antimicrobial pathways, including IL-1β secretion [42]. |
| **CCL3** | C-C motif chemokine ligand 3 (CCL3); aka macrophage inflammatory protein 1-alpha (MIPα); binds to receptor CCR5. | • Chemoattractant for neutrophils, monocytes, and macrophages. All CCR5 ligands (CCL3, CCL4, & CCL5) are upregulated in the lungs of a murine model of *MTB* infection [50]. Yet, the CCR5 knockout mice recruit adequate number of lymphocytes to the lungs and were not compromised in controlling *M. tuberculosis* infection [51]. Infection of mice with the rough strain of *M. avium* (compared to the smooth strain) resulted in more severe disease and greater production of pro-inflammatory cytokines, CCL3, and CCL5 [52]. |
| **TNF** | Tumor necrosis factor | • TNF is critical in host defense against *M. tuberculosis*. It is essential for effective granuloma formation [53].<br>• TNF works synergistically with IFNγ to activate macrophages, enhancing phagolysosome fusion and intracellular killing of mycobacteria [53].<br>• Sustained TNF activity is required to maintain latent tuberculosis. Disruption of TNF activity leads to granuloma breakdown and reactivation of latent infection [54]. |
| **CSF1** | Monocyte colony stimulating factor (M-CSF) | • Repletion during *M. tuberculosis* infection in mice decreases foamy macrophage numbers, increases class 2 MHC molecules on alveolar macrophages, and increases expression of the pattern-recognition receptor DEC-205 [55].<br>• Mobilizes myeloid cells into the blood and tissue and maintains tissue integrity [56]. |
|  |  | • M-CSF activated macrophages skew towards M2 anti-inflammatory phenotype [46, 47, 57], which secrete anti-inflammatory mediators such as IL-4, IL-10, and TGFβ [57]. Activation of M2 macrophages by M-CSF with *M. tuberculosis* infection may protect against excessive inflammation and thus preserve lung function [58].<br>• Macrophages differentiated by M-CSF have increased phagocytosis (due to increased expression of phagocytic receptors) and decreased killing of *M. marinum* than those differentiated by IL-34 [59]. |
| **Selected MAC-downregulated genes** |  |  |
| **MMP15,**<br>**MMP16,**<br>**MMP17** | Matrix metalloproteinases | • MMP15, MMP16, and especially MMP17 are known to cleave *M. tuberculosis* Hsp65 (aka GroEL2) *in vitro*, resulting in digested peptides that are antigenic [60]. Hsp65 is present in all mycobacteria [61].<br>• Full length GroEL2 induces cell surface-associated costimulatory molecules and increases IFNγ, IL-2, and IL-17 from antigen-specific CD4<sup>+</sup> T cells [62]. |
| **CX3CR1** | CX3C chemokine receptor 1 | • Monocytes of individuals with latent tuberculosis infection have increased expression of CX3CR1 on monocytes cell surface [63].<br>• In mice, CX3CR1 deficiency is associated with higher *M. tuberculosis* burden in the mediastinal lymph nodes, making it host protective [64]. However, deficiency of CX3CR1 has no substantial impact on *M. tuberculosis* control in the lungs of mice [64, 65]. |

**aka**=also known as; **IL-1Ra**=interleukin-1 receptor antagonist; **IL-21R**=interleukin 21 receptor; **MAC**=*Mycobacterium avium* complex; **MAIT**=mucosa-associated invariant T cells; **NTM**=non-tuberculous mycobacteria; **PBMC**=peripheral blood mononuclear cells

### Genes regulated by AAT of *MAC*-infected THP-1^control^ macrophages

To determine the effects of AAT in the context of *MAC* infection, bulk RNA sequencing was performed on THP-1^control^ cells infected with *MAC* (THP-1^control^ + *MAC*) as well as THP-1^control^ cells infected with *MAC* and incubated with AAT (THP-1^control^ + *MAC* + AAT). Gene ontology (GO) enrichment analysis was next used to categorize genes upregulated (**Supplemental Figure 2A**) or downregulated (**Supplemental Figure 2B**) by AAT in *MAC*-infected THP-1^control^ cells into various functional categories/pathways. The heat map of specific genes upregulated or downregulated by AAT in *MAC*-infected THP-1^control^ cells were generated. Comparison of *MAC*-infected THP-1^control^ cells ± AAT incubation revealed that 1,200 genes were highly upregulated by AAT; some of these AAT-upregulated genes in the *MAC*-infected cells included *BAX*, *BAD*, and *IL-23A*, (**Supplemental Figure 3**). Compared to *MAC* infection alone, 890 genes were highly downregulated by AAT in the *MAC*-infected THP-1^control^ cells and included IKBKG, *TNF*, *IFNAR1*, *CCL2*, *CSF1*, *IRF3, AKT2*, and *IRF7* (**Supplemental Figure 4**).

*MAC* induced *CSF1* expression and AAT modestly but significantly attenuated this induction (**Figure 3A**). Additionally, AAT inhibition of *MAC*-induced *CSF1* mRNA levels in THP-1^control^ cells was attenuated in THP-1^GR-KD^ cells. To validate this further, we quantified the mRNA of the *CSF1* by RT-qPCR in THP-1^control^ and THP-1^GR-KD^ cells and found that the inhibitory effect of AAT on *CSF1* gene expression is dependent on AAT binding to the GR (**Figure 3B**). However, AAT inhibition of *MAC*-induced M-CSF *protein* was not dependent on GR (**Figure 3C**).

**Figure 3.**
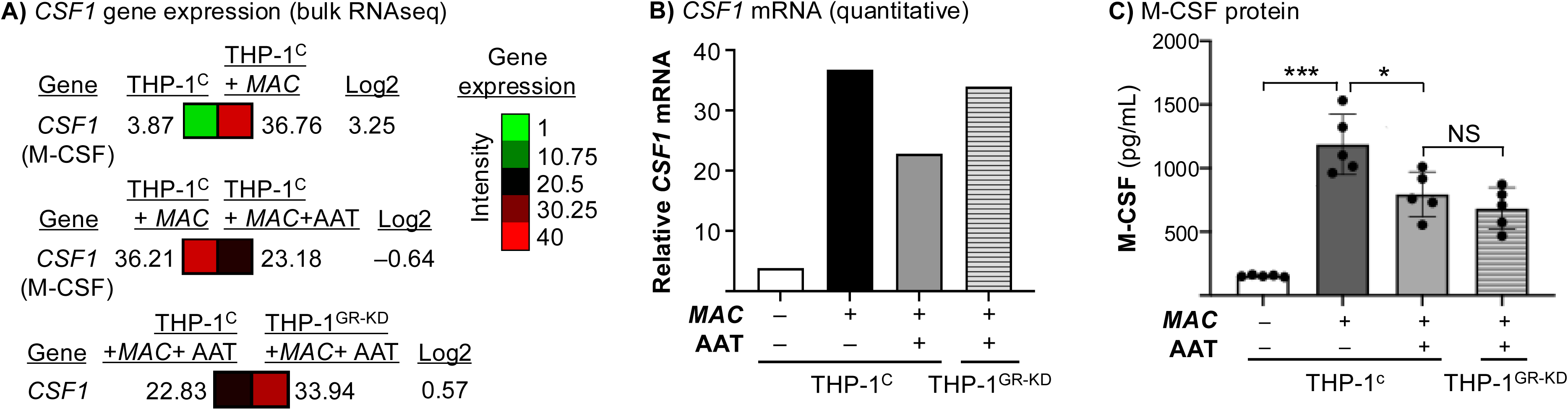
Analysis *CSF1* gene expression regulated by *MAC* infection, AAT, and GR. **(A)** *MAC* significantly induced *CSF1* gene expression in THP-1^C^ cells, defined as ≥ two-fold (log_2_≥1) increase in Reads per Kilobase Million (RPKM) (**top heat map**) and AAT inhibited this induction (**middle heatmap**). The numbers listed below log_2_ were determined by calculating log_2_ of [THP-1^C^+*MAC* divided by THP-1^C^] or THP-1^C^+*MAC*+AAT divided by THP-1^C^+*MAC*. Compared to THP-1C cells incubated with both *MAC*+AAT, THP-1GR-KD cells incubated with *MAC*+AAT showed increase CSF1 gene expression, indicating that AAT inhibited MAC-induced CSF1 gene expression via GR (**bottom heat map)**. The Legend colors for the “*CSF1* gene expression” of the heatmaps were harmonized for direct comparisons. Therefore, the heatmap colors differ from Figure 1A, Supplemental Figure 4B, and Supplemental Figure 6A for *CSF1* but the RPKM values remain the same. **(B)** Relative levels of CSF1 mRNA were quantified via RT-qPCR in THP-1^C^ and THP-1^GR-KD^ incubated with *MAC* ± AAT. **(C)** M-CSF protein was quantified in the supernatant of differentiated THP-1^C^ and THP-1^GR-KD^ cell cultures after incubation with *MAC* ± AAT for three days. Differentiated THP-1^c^ macrophages were left unstimulated or infected with *MAC* ± AAT (bars 1-3). Differentiated THP-1^GR-KD^ macrophages were also incubated with *MAC* + AAT to determine whether GR plays a role in AAT inhibition of *MAC*-induced M-CSF (bar 4).

### AAT-regulated, GR-dependent genes in the context of *MAC* infection

We next determined the role AAT–GR binding may play by comparing the effects of AAT on *MAC*-infected THP-1^control^ and THP-1^GR-KD^ cells. Gene ontology (GO) enrichment analysis was next used to categorize genes upregulated (**Supplemental Figure 5A**) or downregulated (**Supplemental Figure 5B**) by AAT that are GR-dependent in *MAC*-infected THP-1^control^ cells into various functional categories/pathways. Because we cannot be certain that *MAC* does not affect AAT binding to GR, we have chosen to describe the 1,624 genes that are regulated by AAT+*MAC* in THP-1^control^ and enhanced in THP-1^GR-KD^ cells to be genes regulated by AAT+*MAC* and attenuated by GR. Some of these AAT+*MAC* regulated genes that were shown to be attenuated by GR (=enhancement with GR knocked down) include *NOS2*, *IL-12A*, *CCL1*, *CCL2*, *IL6*, and *CSF1* (**Supplemental Figure 6**). Since *MAC*-induced *CSF1* is inhibited by AAT (**Supplemental Figure 4**) and AAT alone did not induce *CSF1* [18], it is reasonable to conclude that AAT inhibition of *MAC*-induced *CSF1* expression is significantly dependent on GR. Conversely, the regulation of 1,683 genes by AAT+*MAC* in THP-1^control^ cells was attenuated in the THP-1^GR-KD^ cells, indicating that the regulation of these genes by AAT+MAC is enhanced by GR. Some of these AAT+*MAC* regulated genes that were shown to be augmented by GR (=attenuation with GR knocked down) include *HLA-DQB1*, *CD80*, *TGFB2*, *IL-23A*, *MMP9*, *IL27*, *TLR2*, and *IL1B* (**Supplemental Figure 7**).

### Biological significance of AAT-regulated GM-CSF with MAC infection

We previously found that AAT induced *CSF2* mRNA expression by 20-fold and GM-CSF protein by > two-fold [18]. In current work, we also found that *MAC* induced *CSF2* gene expression (**Figure 4A**). Since GM-CSF has functions independent of its role in cellular differentiation [42, 48, 49, 66], we examined the effects of neutralizing GM-CSF induced by *MAC* ± AAT in the control of *MAC* infection in THP-1 cells that were differentiated with PMA. The THP-1 cells were infected with *MAC* (MOI 5:1) for one hour, two days, or four days alone or with AAT (3 mg/mL) ± anti-GM-CSF antibody (0.1 and 0.5 µg/mL). Quantitation of *MAC* showed that AAT reduced the cell-associated *MAC* by ∼50% at four days after infection and this reduction was significantly abrogated by anti-GM-CSF antibody (**Figure 4B**).

**Figure 4.**
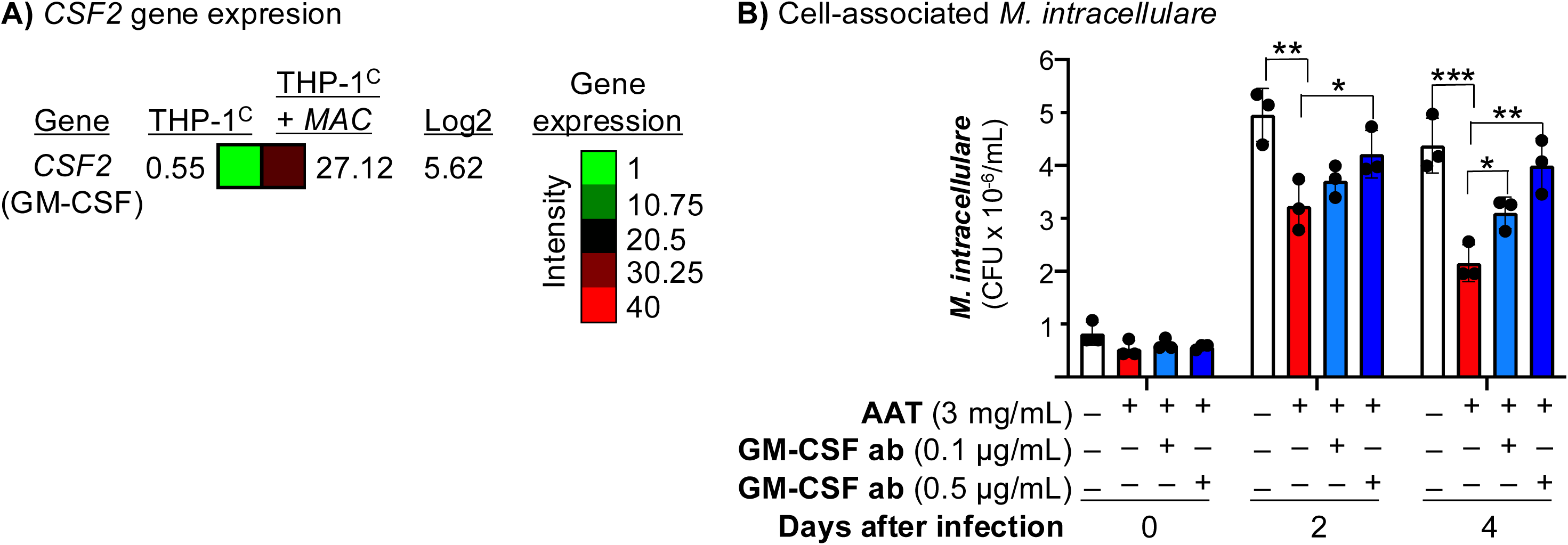
Biologic activity of *MAC*- and/or AAT-induced CSF2 (GM-CSF) against *MAC* in macrophages. **(A)** *MAC* significantly induced *CSF2* gene expression in THP-1^C^ cells by 49-fold, defined as ≥ two-fold (log_2_≥1) increase in Reads per Kilobase Million (RPKM). We previously showed that AAT also induced *CSF2* gene expression by 20-fold and GM-CSF protein expression by >two-fold [18]. **(B)** THP-1 macrophages were infected with *MAC* alone or with AAT, the latter with or without neutralizing antibody to GM-CSF (R&D systems, Cat# MAB215-100). After 1 hour, 2 and 4 days of infection, cell-associated *MAC* were quantified. Data shown are mean ± SEM of three independent experiments with each performed in duplicate wells. *p<0.05, **p<0.01, ***p<0.001. **AAT**=alpha-1-antitrypsin; ***MAC***= *Mycobacterium avium* complex**; THP-1^C^**=control THP-1 cells

## DISCUSSION

Whereas the canonical function of AAT is inhibition of elastase and other serine proteases, AAT also has other functions including mitigation against excessive and injurious inflammation [13–17] and augmentation of host-protective immunity against various pathogens [9, 14, 67–80]. The mechanisms by which AAT antagonizes various pathogens include: *(i)* preventing elastase from cleaving Fcy receptor-1 and complement receptor-1 from cell surface (receptors needed for opsonization and phagocytosis of mycobacreria); *(ii)* inducing autophagy (a mechanism to kill intracellular pathogens); *(iii)* inhibiting entry of SARS-CoV-2 and hepatitis C virus into host cells by inhibiting the serpin TMPRSS2 that processes the surface viral proteins necessary for the intracellular entry of the viral genome; *(iv)* preventing excessive inflammation, mucus hypersecretion, ciliary dysfunction and structural lung injury such as emphysema and bronchiectasis that predispose to infections; and *(v)* mitigating elastase-inhibition of efferocytosis [13, 14, 16, 18, 81–88]. While some of these functions can be attributed to gene regulation by AAT, this area remains poorly investigated.

We previously reported that AAT binds to the transcriptional regulator GR forming the AAT–GR complex, analogous to the glucocorticoid–GR complex. The AAT–GR complex also translocates to the nucleus where it regulates the expression of a panoply of genes involved in inflammation and host immunity [14, 18]. Specifically, AAT–GR complex alone induced the expression of cytokines (IL-1β, IL-23, GM-CSF) [18] that are host-protective against mycobacteria [13, 18, 45, 89, 90]. Therefore, we investigated the gene-regulating effects of AAT–GR in the context of *MAC*-infected macrophages.

Herein, we found that *MAC* induced mRNA expression of many genes including *CSF1* (M-CSF) and *CSF2* (GM-CSF) (**Figure 5A**). Whereas AAT inhibited *MAC*-induced M-CSF mRNA expression, AAT did not inhibit *MAC*-induced GM-CSF mRNA expression. Furthermore, AAT inhibition of *MAC*-induced *CSF1* mRNA was GR-dependent but this GR dependence was lost when CSF-1 (M-CSF) protein was quantified; this finding suggests that AAT also inhibits M-CSF protein expression by a GR-independent mechanism.

**Figure 5.**
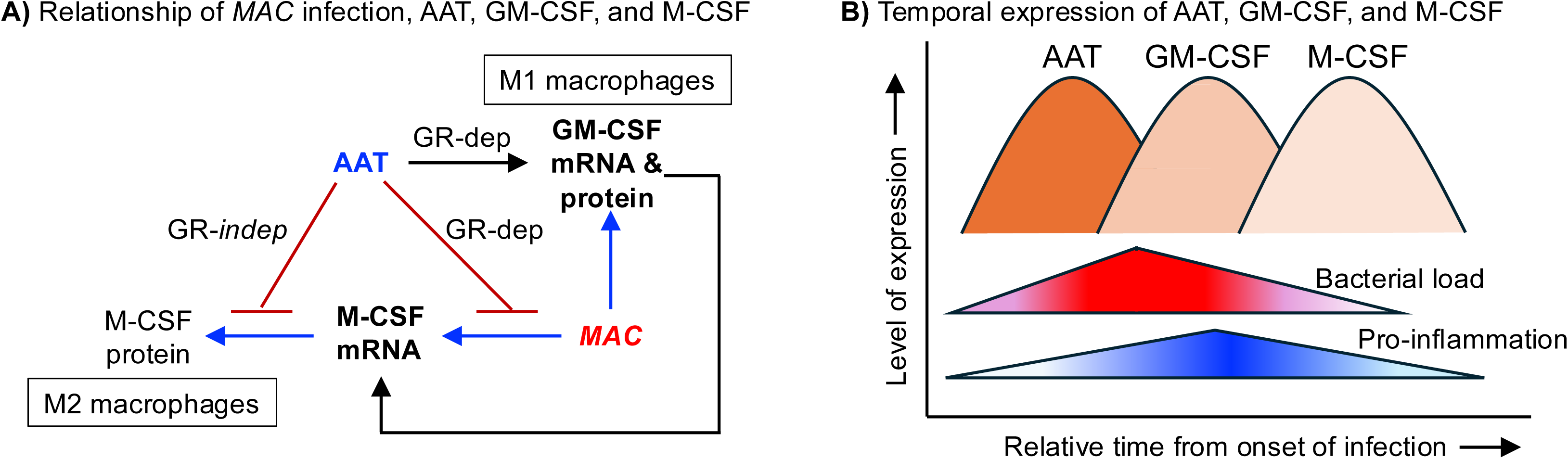
Diagrams showing the relationship between *MAC* infection and AAT with M-CSF and GM-CSF expression. **(A)** Diagram showing the differential effects of AAT on GM-CSF and M-CSF expression. Prior work showed AAT induces GM-CSF expression [18]. *MAC* induces both GM-CSF and M-CSF expression. While AAT inhibits *MAC*-induced M-CSF mRNA expression in a GR-dependent fashion, this dependency was lost when M-CSF protein was quantified. AAT did not inhibit *MAC*-induced GM-CSF expression. **(B)** Hypothesized diagram of the temporal expression of AAT, GM-CSF, and M-CSF. Both *MAC* and AAT induce GM-CSF which helps control the infection by skewing macrophages toward the M1 phenotype and an initial effective inflammatory response to kill mycobacteria; *i.e.*, host-protective inflammation coincides with or soon after peak of GM-CSF expression with reduction of the bacterial load. AAT also inhibits MAC-induced M-CSF to limit M2 macrophages during the initial stage of the infection. Once the infection is under control, AAT decreases and there is increase in M-CSF (from removal of AAT-inhibition of M-CSF expression as well as GM-CSF induction of M-CSF), an anti-inflammatory (“wound healing”) response occurs to blunt host tissue injury. **AAT**=alpha-1-antitrypsin; ***MAC***= *Mycobacterium avium* complex; **M-CSF**=monocyte colony stimulating factor.

GM-CSF skews monocytes towards a pro-inflammatory M1 phenotype whereas M-CSF differentiates macrophages toward the anti-inflammatory M2 phenotype (**Figure 5A**) [41, 43, 46, 47]. A denouement of GM-CSF-induced bias is that the M1 macrophages have greater capacity to control an *M. tuberculosis* infection by a panoply of mechanisms including greater macrophage survival, a stronger *in vitro* granuloma-like response, increased production of IL-1ý, IL-12, and IL-10, decreased excessive levels of TNF and IL-6, increased autophagy, increased phagosome-lysosome fusion, and increased nitric oxide production [42, 44, 91–93]. GM-CSF produced by invariant natural killer T (iNKT) cells also enhanced macrophage control of *M. tuberculosis* infection [48]. Neutralization of GM-CSF in mice promoted M2 macrophage expression, reduced nitric oxide, and higher *M. bovis* BCG burden [91]. Similarly, mice with genetic disruption for GM-CSF were more susceptible to *M. tuberculosis* [94]. GM-CSF was also shown to protect mice against *M. abscessus* infection [45] and to be a promising adjunctive agent for NTM lung disease in humans [95, 96].

In contrast to the effects of GM-CSF, macrophages differentiated with M-CSF and infected with *M. tuberculosis* were not as efficient at activating autologous memory CD4^+^ T cells compared to macrophages that were differentiated with GM-CSF [58]. However, repletion of M-CSF three weeks after *M. tuberculosis* infection in mice led to a decrease in foamy macrophage numbers, increased class 2 MHC molecules on alveolar macrophages, and increased expression of the pattern-recognition receptor DEC-205, suggesting that M-CSF also plays a part in host-defense against mycobacteria [55]. Another host-beneficial function of M-CSF is the mobilization of myeloid cells from the bone marrow into the blood and tissue as well as maintenance of tissue integrity [56]. Additionally, GM-CSF induces M-CSF production in monocytes and macrophages, suggesting that sequential expression of these cytokines may be optimal for host-defense against mycobacterial infections [91, 97]. Thus, it is likely that the relative amounts of GM-CSF and M-CSF produced in a coordinated temporal fashion dictate whether there is optimal innate immune control of infections.

Overall, these results combined with other findings we previously reported suggest that AAT confers host-protection during *MAC* infection by several mechanisms including: *(i)* induction of autophagy of *MAC*-infected cells [13], *(ii)* production of GM-CSF [18], and *(iii)* attenuation of *MAC*-induced M-CSF but not of *MAC*-induced GM-CSF. Therefore, the net effect of AAT during *MAC* infection is increased levels of protective M1 macrophages and decreased levels of less-protective M2 macrophages during the initial stage of the infection. Since AAT is an acute phase reactant, perhaps an early AAT-induced GM-CSF response – with a concomitant AAT inhibition of M-CSF – optimizes control of the initial infection whereas in the resolution phase of the infection, a reduction of AAT to homeostatic levels with less inhibition of M-CSF promotes an anti-inflammatory macrophage phenotype that minimizes injury to host tissues (**Figure 5B**). These findings help to provide a better understanding of host immunity against NTM infection, which, in turn, may lead to the development of more effective treatments.

A limitation to this study is that differentiated THP-1 cells were used and thus the findings are unlikely to be applicable across all human subjects based on genotypic differences. However, this same limitation is also a strength because by employing THP-1 cells, we have been able to create stable populations of THP-1^control^ and THP-1^GR-KD^ cells, resulting in much greater transfection efficiency of the lentivirus-shRNA targeting GR than the strategy of transient GR-knockdown that would have to be employed in primary macrophages. Furthermore, since we wish to compare control cells with cells knocked down for GR, using the THP-1 cell line keeps other variables to a minimum as primary macrophages, even obtained from the same individual, are more likely to have chronotropic fluctuations of their immunologic phenotype. In addition, we and others have shown that the THP-1 cell line mimics qualitatively the responses of primary human macrophages to different stimuli, particularly with mycobacterial infections [98–103].

## Acknowledgments

We thank Miles Pufall, Ph.D. (University of Iowa Carver College of Medicine) for his kind gift of the glucocorticoid receptor (NR3C1) human shRNA (shRNA-GR) Lentiviral Particle. We thank Hua Huang, MD, PhD and Robert Sandhaus, MD, PhD for intellectual input. We are also grateful to Nina C. Khoo, M.D. for her kind support.

## Data Availability Statement

Raw data are available for review.

