## Supplemental Figures and Legend for "Alpha-1-antitrypsin (AAT) inhibits *Mycobacterium intracellulare* induction of monocyte colony stimulating factor: another host-defense function of AAT"

### Supplemental Figure Legend

**Supplemental Figure 1. GO-analysis of *MAC*-regulated genes in THP-1$^{C}$ macrophages. (A)** *MAC*-induced genes sorted by various functional and subcellular categories. The p values were calculated by a one-side Fisher's exact test with the adjustment of Benjamini-Hochberg Method. **(B)** *MAC*-inhibited genes sorted by various functional and subcellular categories. Data represent duplicate of two biological samples analyzed independently. ***MAC***=*Mycobacterium avium* complex; **P II**=polymerase II; **RT**=regulation of transcription; **THP-1$^{C}$**=control THP-1 cells.

**Supplemental Figure 2. GO-analysis of AAT regulated genes in *MAC*-infected THP-1$^{C}$ macrophages. (A)** AAT-induced genes in *MAC*-infected THP-1$^{C}$ cells sorted by various functional and subcellular categories. **(B)** AAT-inhibited genes in *MAC*-infected THP-1$^{C}$ cells sorted by various functional and subcellular categories. The p values were calculated by a one-side Fisher's exact test with the adjustment of Benjamini-Hochberg Method. Data represent duplicate of two biological samples. **AAT**=alpha-1-antitrypsin; **ATP**=adenosine triphosphate; ***MAC***=*Mycobacterium avium* complex.

**Supplemental Figure 3. Heatmap of selected genes upregulated by AAT in *MAC*-infected THP-1$^{C}$ macrophages.** Gene expression in *MAC*-infected THP-1$^{C}$ cells ± AAT was compared to determine what genes are induced by AAT in the context of *MAC* infection. Highly upregulated genes are signified by a change of ≥ 2-fold. The values of THP-1$^{C}$+*MAC* and THP-1$^{C}$+*MAC*+AAT indicate Reads per Kilobase Million (RPKM). The numbers listed below $\log_2$ were determined by calculating $\log_2$ of THP-1$^{C}$+*MAC*+AAT divided by THP-1$^{C}$+*MAC*. Relevant genes were selected and sorted into the following functional categories: **(A)** Autophagy; **(B)** Apoptosis; **(C)** Lysosomes; **(D)** Inflammatory response; and **(E)** Phagocytosis. **AAT**=alpha-1-antitrypsin; **THP-1$^{C}$**=control THP-1 cells; ***MAC***=*Mycobacterium avium* complex.

**Supplemental Figure 4. Heatmap of selected genes downregulated by AAT genes in *MAC*-infected THP-1$^{C}$ macrophages**. Gene expression in *MAC*-infected THP-1$^{C}$ cells ± AAT was compared to determine what genes are inhibited by AAT in the context of *MAC* infection. Highly downregulated genes are signified by a change of ≤ 0.5-fold. The values of THP-1$^{C}$+*MAC* and THP-1$^{C}$+*MAC*+AAT indicate Reads per Kilobase Million (RPKM). The numbers listed below $\log_2$ were determined by calculating $\log_2$ of THP-1$^{C}$+*MAC*+AAT divided by THP-1$^{C}$+*MAC*. Relevant genes were selected and sorted into the following functional categories: **(A)** TLR signaling pathway; **(B)** TNF signaling pathway; **(C)** TLR4 signaling pathway; **(D)** Apoptotic signaling pathway; **(E)** mTOR signaling pathway; and **(F)** Autophagy. **AAT**=alpha-1-antitrypsin; ***MAC***=*Mycobacterium avium* complex; **mTOR**=mammalian target of rapamycin; **THP-1$^{C}$**=control THP-1 cells; **TLR**=toll-like receptor; **TNF**=tumor necrosis factor.

**Supplemental Figure 5. GO-analysis of selected AAT-regulated, GR-dependent genes in *MAC*-infected THP-1 cells. (A)** *MAC*+AAT-regulated genes upregulated by GR sorted by various functional and subcellular categories. **(B)** *MAC*+AAT-regulated genes downregulated by GR sorted by various functional and subcellular categories. The p values were calculated by a one-side Fisher's exact test with the adjustment of Benjamini-Hochberg Method. Data represent duplicate of two biological samples. **AAT**=alpha-1-antitrypsin; **ATP**=adenosine triphosphate; **ErK**=extracellular signal-regulated kinase; **GR**=glucocorticoid receptor; **Mapk**=mitogen-activated protein kinase; **PI3K**=phosphoinositide 3-kinase.

**Supplemental Figure 6. Heatmap of *MAC*+AAT regulated genes that are inhibited by GR.** Gene expression in THP-$1^C$ and THP-$1^{GR\text{-}KD}$ cells (both infected with *MAC* and incubated with AAT) were compared to determine the effect of the GR on AAT gene expression in the context of *MAC* infection. Highly upregulated genes are signified by a change of ≥ 2-fold. The values of THP-$1^C$+*MAC*+AAT and THP-$1^{GR\text{-}KD}$+*MAC*+AAT indicate Reads per Kilobase Million (RPKM). The numbers listed below $\log_2$ were determined by calculating $\log_2$ of THP-$1^{GR\text{-}KD}$+*MAC*+AAT divided by THP-$1^C$+*MAC*+AAT. Relevant genes were selected that were inhibited by AAT in *MAC*-infected THP-$1^C$ cells but induced in the absence of the GR. These genes were sorted into the following functional categories: **(A)** Cytokines; **(B)** Inflammatory response; **(C)** Innate immunity; **(D)** Antiviral defense; **(E)** Tuberculosis; and **(F)** mTOR signaling pathway. **GR**=glucocorticoid receptor; ***MAC***=*Mycobacterium avium* complex; **mTOR**=mechanistic target of rapamycin; **THP-$1^C$**=control THP-1 cells; **THP-$1^{GR\text{-}KD}$**=THP-1 cells knocked down for the glucocorticoid receptor.

**Supplemental Figure 7. Heatmap of *MAC*+AAT regulated genes that are augmented by GR.** Gene expression in THP-$1^C$ and THP-$1^{GR\text{-}KD}$ cells (both infected with *MAC* and incubated with AAT) were compared to determine the effect of the GR on AAT-regulated gene expression in the context of *MAC* infection. Highly downregulated genes are signified by a change of ≤ 0.5-fold. The values of THP-$1^C$+*MAC*+AAT and THP-$1^{GR\text{-}KD}$+*MAC*+AAT indicate Reads per Kilobase Million (RPKM). The numbers listed below $\log_2$ were determined by calculating $\log_2$ of THP-$1^{GR\text{-}KD}$+*MAC*+AAT divided by THP-$1^C$+*MAC*+AAT. Relevant genes were selected that were induced by AAT in MAC-infected THP-$1^C$ cells but inhibited in the absence of the GR. These genes were sorted into the following functional categories: **(A)** Cytokines and chemokines; **(B)** Regulator molecules; **(C)** Protein kinase, and **(D)** Toll-like receptor; **(E)** Apoptosis and autophagy. ***MAC***=*Mycobacterium avium* complex; **THP-$1^C$**=control THP-1 cells; **THP-$1^{GR\text{-}KD}$**=THP-1 cells knocked down for the glucocorticoid receptor.

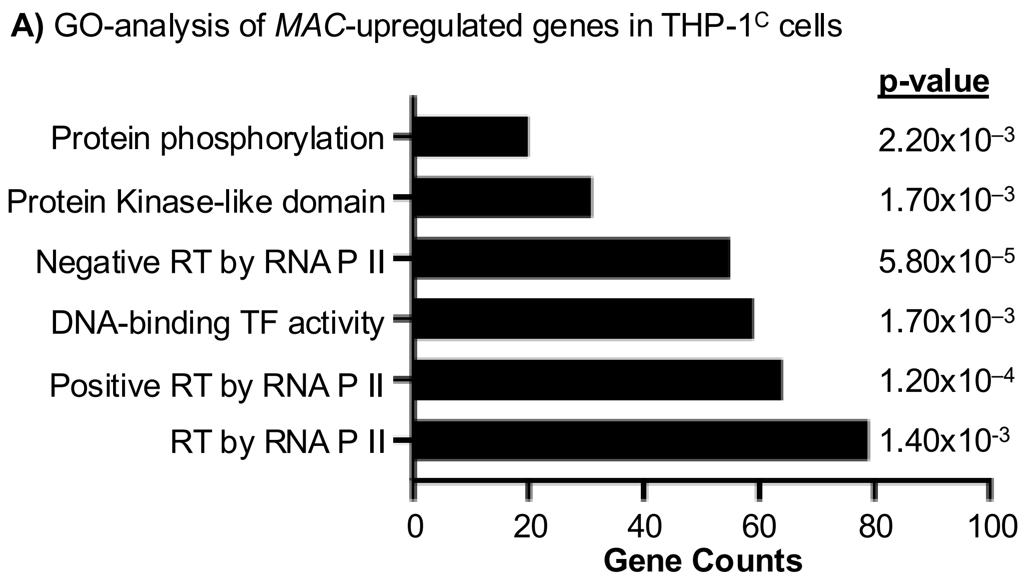


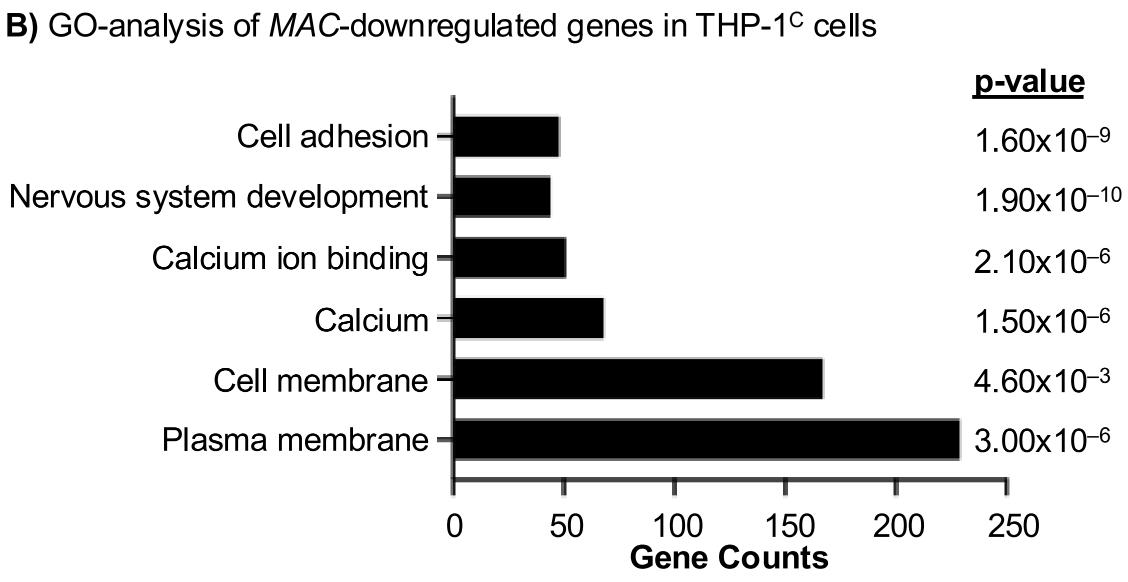


**Supplemental Figure 1**
Bai X, Narum DE et al

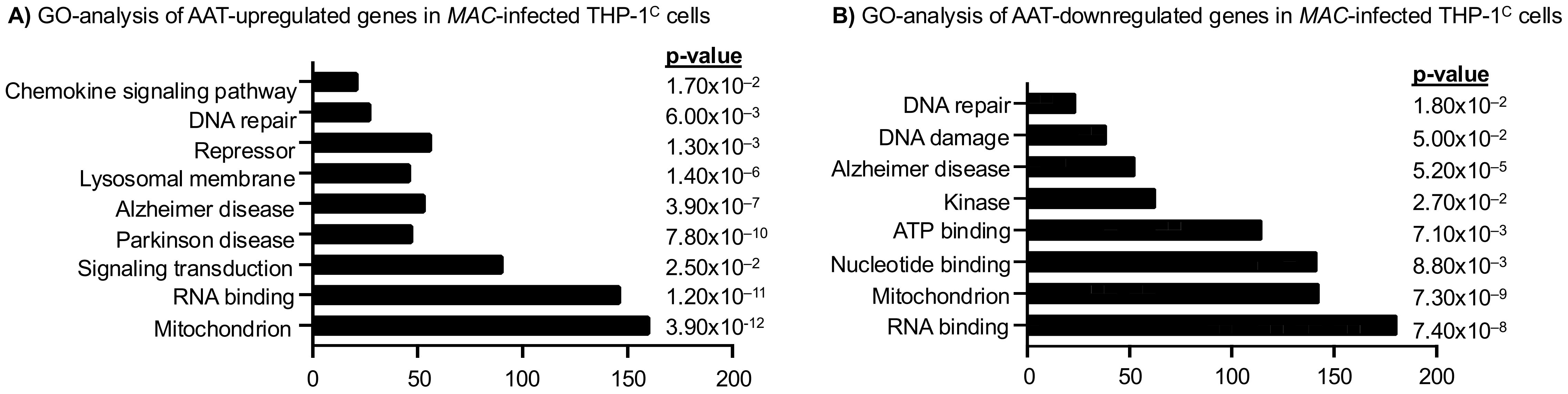


**Supplemental Figure 2**

Bai X, Narum DE et al

**A)** Autophagy

| Genes | THP-1C + *MAC* | THP-1C + *MAC*+AAT | Log2 |
|---|---|---|---|
| MAP1LC3B | 10.80 | 19.85 | 0.88 |
| DEPP1 | 10.10 | 17.19 | 0.77 |
| TBC1D17 | 127.59 | 180.46 | 0.50 |
| ATG9B | 29.57 | 38.19 | 0.37 |
| RNF41 | 15.10 | 18.97 | 0.33 |
| GABARAPL2 | 12.23 | 14.42 | 0.24 |
| TOLLIP | 21.44 | 25.13 | 0.23 |
| ATG4B | 18.39 | 21.22 | 0.21 |
| ATG4D | 36.58 | 41.70 | 0.19 |
| WIPI1 | 11.10 | 12.08 | 0.12 |
| UBQLN1 | 10.35 | 11.22 | 0.12 |
| ATG13 | 11.19 | 12.13 | 0.12 |
| WIPI2 | 10.82 | 11.55 | 0.09 |
| SIRT2 | 14.09 | 15.00 | 0.09 |

**B)** Apoptosis

| Genes | THP-1C + *MAC* | THP-1C + *MAC*+AAT | Log2 |
|---|---|---|---|
| CHAC1 | 11.76 | 32.56 | 1.47 |
| BAX | 13.93 | 24.65 | 0.82 |
| VDAC1 | 25.18 | 38.10 | 0.60 |
| NLRP3 | 11.95 | 16.89 | 0.50 |
| DAD1 | 35.39 | 50.01 | 0.50 |
| DPF2 | 10.08 | 14.04 | 0.48 |
| NFKB1 | 11.06 | 15.30 | 0.47 |
| PHLDA2 | 64.20 | 88.22 | 0.46 |
| FIS1 | 15.19 | 20.61 | 0.44 |
| PPID | 28.01 | 37.80 | 0.43 |

**B)** Apoptosis (continued)

| Genes | THP-1C + *MAC* | THP-1C + *MAC*+AAT | Log2 |
|---|---|---|---|
| TCIRG1 | 226.33 | 302.66 | 0.42 |
| MEF2C | 16.98 | 22.32 | 0.39 |
| MFSD10 | 26.27 | 34.14 | 0.38 |
| CHST11 | 117.31 | 149.23 | 0.35 |
| STK17A | 14.12 | 17.63 | 0.32 |
| PLK3 | 22.92 | 27.96 | 0.29 |
| MARCKS | 14.22 | 17.33 | 0.29 |
| RNF34 | 97.72 | 117.99 | 0.27 |
| FAM32A | 11.08 | 13.36 | 0.27 |
| BAD | 12.16 | 14.64 | 0.27 |
| BNIPL | 314.52 | 375.94 | 0.26 |
| TCHP | 223.30 | 266.69 | 0.26 |
| PTEN | 16.37 | 19.11 | 0.22 |
| UNC5B | 40.06 | 46.60 | 0.22 |
| BAG6 | 37.00 | 42.11 | 0.19 |
| SOD1 | 52.57 | 59.83 | 0.19 |
| RASSF5 | 16.28 | 18.36 | 0.17 |
| PYCARD | 34.71 | 38.93 | 0.17 |
| INPP5D | 19.85 | 21.95 | 0.15 |
| AIMP2 | 20.35 | 22.40 | 0.14 |
| TRAF7 | 34.45 | 37.43 | 0.12 |
| GHITM | 11.83 | 12.83 | 0.12 |
| PPP1R15A | 36.42 | 39.31 | 0.11 |
| GPX4 | 138.52 | 149.29 | 0.11 |
| BCL7B | 21.12 | 22.65 | 0.10 |
| PACS2 | 15.21 | 16.28 | 0.10 |
| LGALS1 | 529.38 | 565.25 | 0.09 |
| VIL1 | 48.83 | 51.96 | 0.09 |

**C)** Lysosomes

| Genes | THP-1C + *MAC* | THP-1C + *MAC*+AAT | Log2 |
|---|---|---|---|
| AP1S2 | 19.71 | 34.39 | 0.80 |
| GGA1 | 18.40 | 24.22 | 0.40 |
| CLTB | 12.60 | 16.50 | 0.39 |
| ATP6AP1 | 88.83 | 115.96 | 0.38 |
| AP3D1 | 17.86 | 22.89 | 0.36 |
| IGF2R | 10.95 | 12.94 | 0.24 |
| AP1M1 | 11.28 | 12.90 | 0.19 |
| GNPTG | 47.42 | 53.36 | 0.17 |

**D)** Inflammatory response

| Genes | THP-1C + *MAC* | THP-1C + *MAC*+AAT | Log2 |
|---|---|---|---|
| ZNF580 | 18.56 | 40.44 | 1.12 |
| CCL3L1 | 42.59 | 71.84 | 0.75 |
| C5AR1 | 11.86 | 17.56 | 0.57 |
| C3 | 29.63 | 42.50 | 0.52 |
| CCL3 | 124.48 | 172.81 | 0.47 |
| AZU1 | 18.85 | 24.32 | 0.37 |
| IL23A | 753.17 | 970.85 | 0.37 |
| NFKBID | 13.81 | 17.18 | 0.32 |
| MAPKAPK2 | 16.64 | 20.04 | 0.27 |
| PRDX5 | 21.80 | 25.94 | 0.25 |
| CD44 | 389.58 | 438.80 | 0.17 |
| CXCL1 | 12.22 | 13.66 | 0.16 |
| HOOK2 | 1163.54 | 1279.12 | 0.14 |
| ADGRE5 | 100.27 | 109.95 | 0.13 |
| CCL20 | 203.31 | 221.21 | 0.12 |
| C4A | 22.05 | 23.77 | 0.11 |
| TGFB1 | 168.97 | 182.11 | 0.11 |
| LOXL3 | 59.44 | 63.92 | 0.10 |

**E)** Phagocytosis

| Genes | THP-1C + *MAC* | THP-1C + *MAC*+AAT | Log2 |
|---|---|---|---|
| CDC42SE1 | 24.28 | 30.27 | 0.32 |
| CNN2 | 25.50 | 28.25 | 0.15 |
| CORO1A | 142.94 | 154.66 | 0.11 |
| GAS6 | 10.35 | 11.18 | 0.11 |
| ITGAL | 23.05 | 25.89 | 0.17 |
| ITGB1 | 16.65 | 24.87 | 0.58 |
| LYST | 22.38 | 27.07 | 0.27 |
| PIP5K1C | 11.25 | 15.26 | 0.44 |
| PLD4 | 19.12 | 25.15 | 0.40 |

**Gene Expression**

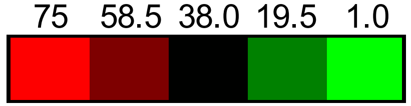


**Supplemental Figure 3**
Bai X, Narum DE et al

### A) TLR signaling pathway

| Genes | THP-1$^{C}$ + *MAC* | THP-1$^{C}$ + *MAC*+AAT | Log2 |
|---|---|---|---|
| IKBKG | 13.19 | 4.95 | -1.42 |
| TNF | 10.75 | 4.21 | -1.35 |
| TRAF3 | 13.19 | 5.46 | -1.27 |
| IFNAR1 | 10.83 | 6.00 | -0.85 |
| FOS | 26.43 | 15.76 | -0.75 |
| TICAM2 | 39.00 | 24.16 | -0.69 |
| IRF3 | 53.19 | 37.57 | -0.50 |
| CCL4L2 | 20.10 | 14.41 | -0.48 |
| SPP1 | 374.88 | 271.14 | -0.47 |
| AKT2 | 15.30 | 11.07 | -0.47 |
| MAP2K3 | 31.92 | 23.64 | -0.43 |
| IRF7 | 42.51 | 33.21 | -0.36 |
| RELA | 34.01 | 26.89 | -0.34 |
| IRAK1 | 31.80 | 25.64 | -0.31 |
| MAPK3 | 24.37 | 20.95 | -0.22 |
| MAP2K7 | 10.05 | 8.69 | -0.21 |
| CCL4 | 41.46 | 36.24 | -0.19 |

**Gene Expression**

75 58.5 38.0 19.5 1.0

### B) TNF signaling pathway

| Genes | THP-1$^{C}$ + *MAC* | THP-1$^{C}$ + *MAC*+AAT | Log2 |
|---|---|---|---|
| CCL2 | 273.59 | 159.70 | -0.78 |
| CSF1 | 36.21 | 23.18 | -0.64 |
| CREB3L2 | 12.12 | 8.51 | -0.51 |
| CXCL2 | 12.48 | 9.41 | -0.41 |
| TNFRSF1A | 22.86 | 17.72 | -0.37 |
| BCL3 | 58.31 | 46.82 | -0.32 |
| BIRC2 | 15.81 | 12.83 | -0.30 |
| RHBDF2 | 69.80 | 57.39 | -0.28 |
| DNM1L | 18.04 | 15.02 | -0.26 |

A red dashed box outlines the CSF1 row.

### C) TLR4 signaling pathway

| Genes | THP-1$^{C}$ + *MAC* | THP-1$^{C}$ + *MAC*+AAT | Log2 |
|---|---|---|---|
| PIK3AP1 | 17.88 | 12.36 | -0.53 |
| RIPK2 | 29.05 | 21.63 | -0.43 |

### D) Apoptotic signaling pathway

| Genes | THP-1$^{C}$ + *MAC* | THP-1$^{C}$ + *MAC*+AAT | Log2 |
|---|---|---|---|
| BCL2L1 | 11.93 | 9.02 | -0.40 |
| PML | 22.28 | 17.25 | -0.37 |
| SOD2-OT1 | 483.92 | 398.65 | -0.28 |
| SOD2 | 483.92 | 398.65 | -0.28 |
| CHEK2 | 20.52 | 17.33 | -0.24 |
| BCL2A1 | 15.90 | 13.90 | -0.19 |
| BCL2L2 | 23.00 | 20.13 | -0.19 |

### E) mTOR signaling pathway

| Genes | THP-1$^{C}$ + *MAC* | THP-1$^{C}$ + *MAC*+AAT | Log2 |
|---|---|---|---|
| EIF4EBP1 | 63.72 | 43.19 | -0.56 |
| EIF4A2 | 30.78 | 23.19 | -0.41 |
| RHEB | 10.78 | 8.35 | -0.37 |
| RPS6 | 307.93 | 261.71 | -0.23 |
| EIF4B | 30.99 | 26.58 | -0.22 |
| EIF4G3 | 14.66 | 12.65 | -0.21 |
| EIF3A | 11.81 | 10.41 | -0.18 |

### F) Autophagy

| Genes | THP-1$^{C}$ + *MAC* | THP-1$^{C}$ + *MAC*+AAT | Log2 |
|---|---|---|---|
| VPS51 | 47.04 | 26.69 | -0.82 |
| TBC1D5 | 15.59 | 10.36 | -0.59 |
| TMBIM6 | 72.41 | 49.53 | -0.55 |
| NPC1 | 16.51 | 11.80 | -0.48 |
| RAB1A | 14.98 | 10.73 | -0.48 |
| VMP1 | 12.05 | 9.25 | -0.38 |
| CALCOCO2 | 11.36 | 8.87 | -0.36 |
| GRAMD1A | 27.03 | 21.52 | -0.33 |
| TRAPPC4 | 21.57 | 17.24 | -0.32 |
| COL6A1 | 52.88 | 42.78 | -0.31 |
| TBC1D25 | 34.86 | 28.77 | -0.28 |
| ATG101 | 13.73 | 12.00 | -0.19 |
| CHMP1A | 12.63 | 11.06 | -0.19 |
| VCP | 26.66 | 23.57 | -0.18 |

**Supplemental Figure 4**

Bai X, Narum DE et al

**A)** GO-analysis of [AAT+*MAC*]-regulated genes upregulated by GR

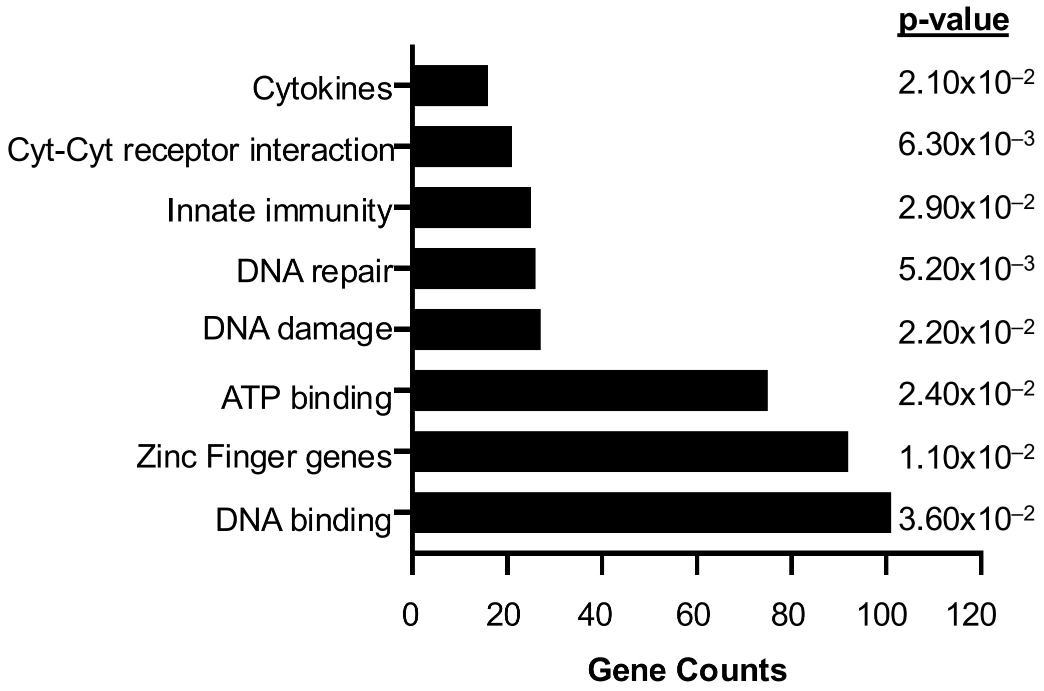


**B)** GO-analysis [AAT+*MAC*]-regulated genes downregulated by GR

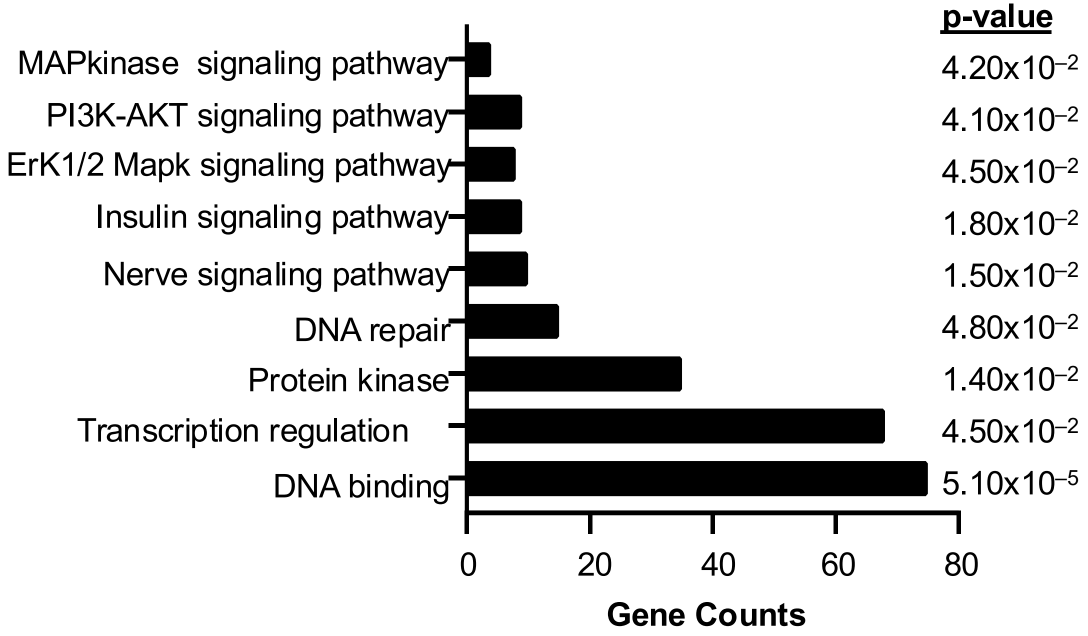


**Supplemental Figure 5**
Bai X, Narum DE et al

### A) Cytokines

| Genes | THP-1$^{C}$ +*MAC*+ AAT | THP-1$^{GR\text{-}KD}$ +*MAC*+ AAT | Log2 |
|---|---|---|---|
| ACVR1C | 0.04 | 0.63 | 3.94 |
| CCR1 | 0.42 | 2.35 | 2.47 |
| TNFSF11 | 0.17 | 0.92 | 2.43 |
| TNFSF13B | 0.37 | 1.92 | 2.39 |
| CXCL6 | 1.02 | 5.25 | 2.36 |
| TNFSF10 | 4.22 | 20.87 | 2.31 |
| ACVR1 | 0.65 | 2.87 | 2.15 |
| CCL13 | 1.47 | 6.29 | 2.09 |
| CSF3 | 0.33 | 1.36 | 2.04 |
| LIF | 0.59 | 2.25 | 1.93 |
| TNFSF4 | 0.54 | 1.96 | 1.86 |
| IL20RB | 2.27 | 8.20 | 1.85 |
| NODAL | 0.40 | 1.42 | 1.84 |
| CXCL3 | 1.20 | 4.11 | 1.78 |
| IL12A | 0.39 | 1.27 | 1.71 |
| GDF1 | 0.77 | 2.42 | 1.66 |
| CCL1 | 181.47 | 567.31 | 1.64 |
| LEPR | 2.10 | 6.11 | 1.54 |
| CCL2 | 157.34 | 447.52 | 1.51 |
| GDF9 | 1.02 | 2.83 | 1.48 |
| IL6 | 1.94 | 5.34 | 1.46 |
| CSF1 | 22.83 | 33.94 | 0.57 |

### B) Inflammatory response

| Genes | THP-1$^{C}$ +*MAC*+ AAT | THP-1$^{GR\text{-}KD}$ +*MAC*+ AAT | Log2 |
|---|---|---|---|
| NOS2 | 0.24 | 1.31 | 2.43 |
| CCRL2 | 1.28 | 6.82 | 2.42 |
| CD14 | 0.61 | 3.20 | 2.40 |
| NOD1 | 0.66 | 3.14 | 2.25 |
| TNFAIP6 | 0.59 | 2.81 | 2.25 |
| GPER1 | 0.15 | 0.54 | 1.84 |
| ORM1 | 0.60 | 2.13 | 1.82 |
| NKG7 | 2.35 | 8.04 | 1.77 |
| NFAM1 | 0.39 | 1.31 | 1.76 |
| DHRS7B | 1.86 | 5.42 | 1.54 |

### C) Innate immunity

| Genes | THP-1$^{C}$ +*MAC*+ AAT | THP-1$^{GR\text{-}KD}$ +*MAC*+ AAT | Log2 |
|---|---|---|---|
| LGR4 | 0.28 | 3.54 | 3.68 |
| RAP1GAP | 0.16 | 1.65 | 3.39 |
| CD86 | 0.16 | 1.15 | 2.80 |
| CFH | 0.18 | 0.84 | 2.20 |
| CFI | 0.69 | 3.02 | 2.13 |
| CD84 | 0.04 | 0.15 | 1.75 |
| VSIG4 | 0.34 | 1.13 | 1.72 |
| TBKBP1 | 1.49 | 4.66 | 1.65 |
| NUDCD1 | 1.15 | 3.45 | 1.58 |
| ERAP2 | 0.21 | 0.62 | 1.58 |

### D) Antiviral defense

| Genes | THP-1$^{C}$ +*MAC*+ AAT | THP-1$^{GR\text{-}KD}$ +*MAC*+ AAT | Log2 |
|---|---|---|---|
| ISG20 | 0.57 | 8.68 | 3.92 |
| IFIT5 | 0.28 | 1.54 | 2.48 |
| IFIT1 | 0.58 | 3.21 | 2.47 |
| IFIT3 | 2.68 | 13.10 | 2.29 |
| MX1 | 0.27 | 1.14 | 2.08 |
| IFIH1 | 2.03 | 7.93 | 1.96 |
| MX2 | 3.70 | 13.56 | 1.87 |
| PARP9 | 1.54 | 5.24 | 1.77 |
| OAS2 | 4.21 | 14.14 | 1.75 |
| DDX60 | 1.45 | 4.66 | 1.68 |
| OASL | 1.50 | 4.65 | 1.63 |
| IFIT2 | 1.25 | 3.78 | 1.60 |
| TRIM22 | 1.26 | 3.68 | 1.54 |

### E) Tuberculosis

| Genes | THP-1$^{C}$ +*MAC*+ AAT | THP-1$^{GR\text{-}KD}$ +*MAC*+ AAT | Log2 |
|---|---|---|---|
| HLA-DRA | 0.35 | 1.89 | 2.42 |
| HLA-DMA | 1.19 | 4.68 | 1.97 |
| HLA-DMB | 1.19 | 4.68 | 1.97 |
| ITGAM | 0.27 | 1.02 | 1.92 |
| SYK | 0.67 | 2.30 | 1.78 |
| FCGR1A | 1.19 | 3.46 | 1.54 |
| CD209 | 2.66 | 7.53 | 1.50 |

### F) mTOR signaling pathway

| Genes | THP-1$^{C}$ +*MAC*+ AAT | THP-1$^{GR\text{-}KD}$ +*MAC*+ AAT | Log2 |
|---|---|---|---|
| FZD5 | 0.05 | 0.23 | 2.20 |
| EIF4E | 1.42 | 6.10 | 2.10 |
| WNT4 | 0.10 | 0.40 | 2.06 |
| IRS1 | 0.22 | 0.84 | 1.93 |
| RPS6KB2 | 5.16 | 15.66 | 1.60 |

**Gene Expression**

10.00 7.75 5.50 3.25 1.00

**Supplemental Figure 6**

Bai X, Narum DE et al

#### A) Cytokines and chemokines

| Genes | THP-1^C + *MAC*+AAT | THP-1^GR-KD + *MAC*+AAT | Log2 |
|---|---|---|---|
| CD80 | 3.13 | 0.40 | -2.97 |
| TGFB2 | 0.21 | 0.05 | -2.10 |
| SOS1 | 3.28 | 0.81 | -2.02 |
| IL-23A | 95.64 | 47.15 | -1.49 |
| ANGPT2 | 0.28 | 0.11 | -1.20 |
| CCL3 | 170.26 | 75.42 | -1.18 |
| CCL3L1 | 70.77 | 34.20 | -1.04 |
| NCF1 | 38.98 | 19.09 | -1.03 |
| IL-27 | 25.10 | 19.50 | -0.91 |
| CCL20 | 217.92 | 117.95 | -0.89 |
| CCL19 | 1.62 | 1.10 | -0.48 |
| IL-1B | 145.13 | 102.87 | -0.47 |

#### B) Regulator molecule

| Genes | THP-1^C + *MAC*+AAT | THP-1^GR-KD + *MAC*+AAT | Log2 |
|---|---|---|---|
| HLA-DQB1 | 2.07 | 0.19 | -3.38 |
| HLA-F | 0.97 | 0.27 | -1.81 |
| SOCS5 | 0.87 | 0.26 | -1.73 |
| ZNF136 | 0.32 | 0.10 | -1.72 |
| NXF3 | 5.13 | 1.57 | -1.71 |
| HLA-G | 3.76 | 1.33 | -1.50 |
| HLA-DRB5 | 2.28 | 0.83 | -1.45 |
| ZNF10 | 5.16 | 1.91 | -1.43 |
| CHUK | 2.23 | 1.06 | -1.08 |
| SRSF8 | 2.64 | 1.30 | -1.02 |

#### C) Protein kinase

| Genes | THP-1^C + *MAC*+AAT | THP-1^GR-KD + *MAC*+AAT | Log2 |
|---|---|---|---|
| PRKCE | 3.33 | 0.34 | -3.29 |
| SRPK3 | 1.83 | 0.19 | -3.28 |
| TTBK2 | 3.57 | 0.40 | -3.15 |
| RIOK2 | 3.03 | 0.45 | -2.76 |
| PRKY | 0.26 | 0.05 | -2.35 |
| TAOK1 | 5.70 | 1.37 | -2.06 |
| WNK4 | 0.27 | 0.07 | -1.97 |
| DYRK2 | 6.01 | 1.56 | -1.95 |
| ATM | 7.32 | 2.00 | -1.87 |
| SGK3 | 5.46 | 1.54 | -1.83 |
| PRKD2 | 14.97 | 5.99 | -1.32 |
| GTF2IP4 | 1.35 | 0.61 | -1.14 |
| CIT | 4.13 | 1.91 | -1.11 |

#### D) Toll like receptor

| Genes | THP-1^C + *MAC*+AAT | THP-1^GR-KD + *MAC*+AAT | Log2 |
|---|---|---|---|
| IL-12RB2 | 0.86 | 0.12 | -2.89 |
| TLR-8 | 1.91 | 0.60 | -1.66 |
| TLR-2 | 8.62 | 5.79 | -0.57 |
| TLR-6 | 2.07 | 1.19 | -0.30 |

#### E) Apoptosis & autophagy

| Genes | THP-1^C + *MAC*+AAT | THP-1^GR-KD + *MAC*+AAT | Log2 |
|---|---|---|---|
| PIK3R4 | 2.91 | 0.38 | -2.94 |
| PIK3R3 | 0.68 | 0.11 | -2.66 |
| ATG4A | 0.91 | 0.18 | -2.36 |
| TGFB2 | 0.20 | 0.05 | -2.1 |
| BAX | 24.23 | 7.42 | -1.71 |
| PIK3R1 | 7.70 | 2.30 | -1.71 |
| TECPR1 | 8.29 | 2.92 | -1.5 |
| ATG14 | 2.59 | 1.02 | -1.34 |
| ATG4C | 3.10 | 1.22 | -1.34 |
| APAF1 | 3.98 | 1.69 | -1.23 |
| TP53INP2 | 6.41 | 2.91 | -1.14 |
| BBC3 | 10.66 | 4.98 | -1.09 |
| MMP9 | 464.10 | 240.74 | -0.96 |
| BCL3 | 46.12 | 36.63 | -0.34 |

**Gene Expression**

10.00 7.75 5.50 3.25 1.00

**Supplemental Figure 7**
Bai X, Narum DE et al
